# Genomic insights into greater yam tuber quality traits

**DOI:** 10.1101/2023.03.17.532727

**Authors:** Ana Zotta Mota, Komivi Dossa, Mathieu Lechaudel, Denis Cornet, Pierre Mournet, David Lopez, Hana Chaïr

## Abstract

Yams are important tubers widely consumed in developing countries in various forms, mainly boiled, pounded and fried. Tuber quality is a key criterion for acceptance by the various actors in the value chain. However, the genetics of yam tuber quality has not yet been comprehensively investigated. Given this lack of knowledge, we used population genomics and candidate gene association approaches to unravel the genetic basis of the three main quality attributes of boiled yam, namely texture, starch content and colour indices. We re-sequenced the whole genome of 127 yam genotypes with different quality attributes, and performed an enrichment of the already available functional genome annotation using comparative genomics. Population genomics analysis highlighted three main genetic groups and 1,053 genes under selection. We focused this work on three known quality trait-related pathways: pectin, starch content and flavonoid biosynthesis, and inventoried in the genome all the structural genes related to those pathways by comparative genomics. We improved the functional annotation of the three pathways with 48% more genes. A total of 18 candidate genes containing genetic variants significantly associated with the target traits were detected, including eight genes that were also found under selection. The texture-related candidate genes were distributed between the pectin and starch pathways. Overall, the use of comparative genomics has allowed the generation of an unprecedented genomic resource. The improved functional annotation of the yam genome is a promising predictive tool for identifying other core genes associated with any trait of interest to meet the growing need and diversified demands for yams.

## INTRODUCTION

Releasing varieties with enhanced quality is a target in the breeding process, and failure to achieve it can lead to the rejection of improved varieties. However, food quality is a complex attribute that reflects the preferences of all actors in the value chain: producers, processors, retailers and consumers. In recent years, there has been an increasing amount of studies on the identification of genomic regions associated with quality traits (Liao et al., 2021; Xia et al., 2020), with a focus on economically important crops such as rice, tomato, citrus, etc. (Xiao et al., 2021; Dominguez et al., 2020; Butelli et al., 2019). Underutilised crops such as tropical root and tuber crops have been largely neglected. However, root and tuber crops are among the most important crops for subsistence and commercial purposes after cereals.

Yams, of the Dioscoreaceae family, are important edible tuber crops, mainly in developing countries. The most widespread species is greater yam (*Dioscorea alata* L.), cultivated mainly for its starchy tubers in tropical regions (Wu et al., 2014; Siqueira et al., 2014; Girma et al., 2014 Lebot et al., 1998). Greater yam displays a wide tolerance to different environments (Orkwor and Asadu, 1998). It is a dioecious and autopolyploid species (2n = 2x = 40, 3x = 60 and 4x = 80) (Arnau et al., 2009). The recent publication of the genetic map of this species (Cormier et al., 2021) followed by the release of its reference genome (Bredeson et al., 2022) have paved the way to work on the genetic architectures of the traits of interest. Genomic regions associated to anthracnose resistance, tuber oxidative browning, tuber morphological attributes, and dry matter have been revealed (Bredeson et al., 2022; Gatarira et al., 2020; Ehounou et al., 2022). Although it is well known that tuber quality is important for varieties adoption in root and tuber crops (Dufour et al. 2021), the genetics of yam tuber quality has not yet been comprehensively investigated.

Greater yam is consumed in various forms, mainly boiled, pounded, fried or baked (Honfozo et al., 2021). The quality of boiled yam, as characterised by sensory testing with consumers, is related to white colour, crumbly, sticky to the fingers, non-fibrous and easy to chew texture, sweet taste and good smell (Honfozo et al., 2021). The choice of colour and the consistency of the tubers after cooking are essential characteristics that have been previously selected in other crops, such as cassava and potato (Alves-Pereira et al., 2022; Hardigan et al., 2017). Texture, which determines the ability of the raw material to soften after cooking while maintaining its firmness, is a complex trait. The main factors influencing texture are starch content and its distribution in the tuber, cell wall structure and composition, and degradation of the middle lamella of the cell wall (reviewed in Taylor et al., 2008; Ross et al., 2011). The cell wall is composed of 90-95% polysaccharides, divided into cellulose, hemicellulose and pectin, and only 5-10% protein. Cell wall polysaccharides and starch content are mainly associated with two metabolic pathways: pentose and glucuronate interconversions, and starch and sucrose metabolism. Colour, one of the most important varietal rejection factors, is partly determined by genes in the flavonoid biosynthetic pathway (He et al., 2021). Despite their importance for the main quality attributes of boiled yam, these pathways have been very little studied.

Over the last decade, hundreds of genomes have been released. Likewise, much efforts have been put on improving the genome annotation in order to extract biological knowledge from genomic sequences (Marks et al., 2021). However, annotating a genome is time and resources consuming. Automatic pipelines can produce inaccurate genome annotation and their results often require manual curation (Bolger et al., 2017). In non-model species, this can be especially true, since several genes lack a precise functional annotation, relying on public databases and gene similarity only. The use of comparative analysis allows to find homologous and orthologous genes from different species and within the species of interest (Mota et al., 2020), and predict more precisely their functions. Using the annotation produced with non-manual curated methods (similarity-based), together with comparative genomics methods can increase the number of genes with a functional annotation, and also help to better assign the functional annotation of other genes.

In this study, we re-sequenced the whole genome of 127 yam genotypes with different quality traits from distinct geographical locations. We focused our work on the three metabolic pathways involved in texture and in starch content (pentose and glucuronate interconversions, starch and sucrose metabolism), and in colour (flavonoid biosynthesis). Through the use of comparative genomics, we were able to improve the annotation of a number of genes, including those in the three major metabolic pathways targeted, thereby enriching the annotation of the *D. alata* genome. We conducted a population genomics study to identify genes under selection. To complement our results, we performed a candidate gene association analysis using phenotypic data on texture, colour and starch content. We identified candidate genes associated with key quality traits for boiled yam and assessed their expression profiles using transcriptomic analysis. Overall, we generated a consistent and valuable amount of genomic resources for future research on important agronomic traits in yam.

## RESULTS

### Pattern of genome-wide variation, population structure and signatures of selection

We generated the whole-genome sequences of 127 genotypes of *Dioscorea* spp. from diverse origins (Africa, Caribbean, Pacific and Asia) (Figure 1A) using short-read sequencing with a mean coverage of 37X (Supplemental Figure 1, Supplemental Table 1). The polymorphism analyses in *D. alata* genotypes yielded a total of 63 M SNPs after filtering. The type of SNPs was divided as 2.8% of variants present in exons, 1.3% of synonymous variants, and 48% intergenic variants (Figure 1B, Supplemental Figure 2). For diversity purposes, only diploid genotypes (107) were used (Supplemental Table 1), since the diversity of the diploid genotypes represents the overall diversity of the species as demonstrated previously (Sharif et al., 2020). Three major groups were separated phylogenetic tree (Supplemental Figure 3A) and by principal component analysis (PCA). A first group consisting mainly of Asian genotypes, a second group composed of Pacific genotypes, and a final group of African and Caribbean genotypes (Figure 1C), which agrees with our previous study, performed by the genotype-by-sequencing analysis on a larger *D. alata* population (Sharif et al., 2020). The genetic structure separated the genotypes into four clusters (Figure 1D). However, as the fourth group had only six genotypes, a robust statistical analysis could not be conducted. Consequently, we conducted our further analysis with *K=3*. The cluster A composed mainly of Pacific representatives and, to a lesser extent, African and Caribbean genotypes, while the cluster B was composed mostly by representatives from Africa and Caribbean, and finally the cluster C had mostly representatives from Asia (26 out of 30) (Figure 1D, Supplemental Figure 3B), which is in accordance with the PCA results. The nucleotide diversity (π) was low in all gene pools 0.407e^-4^. The π values were significantly different between the clusters, with the highest values obtained for the cluster A and C (π = 0.399e^−4^ and 0.396e^−4^, respectively) and the lowest for the Cluster B (π = 0.33e^−4^).

**Figure 1:**
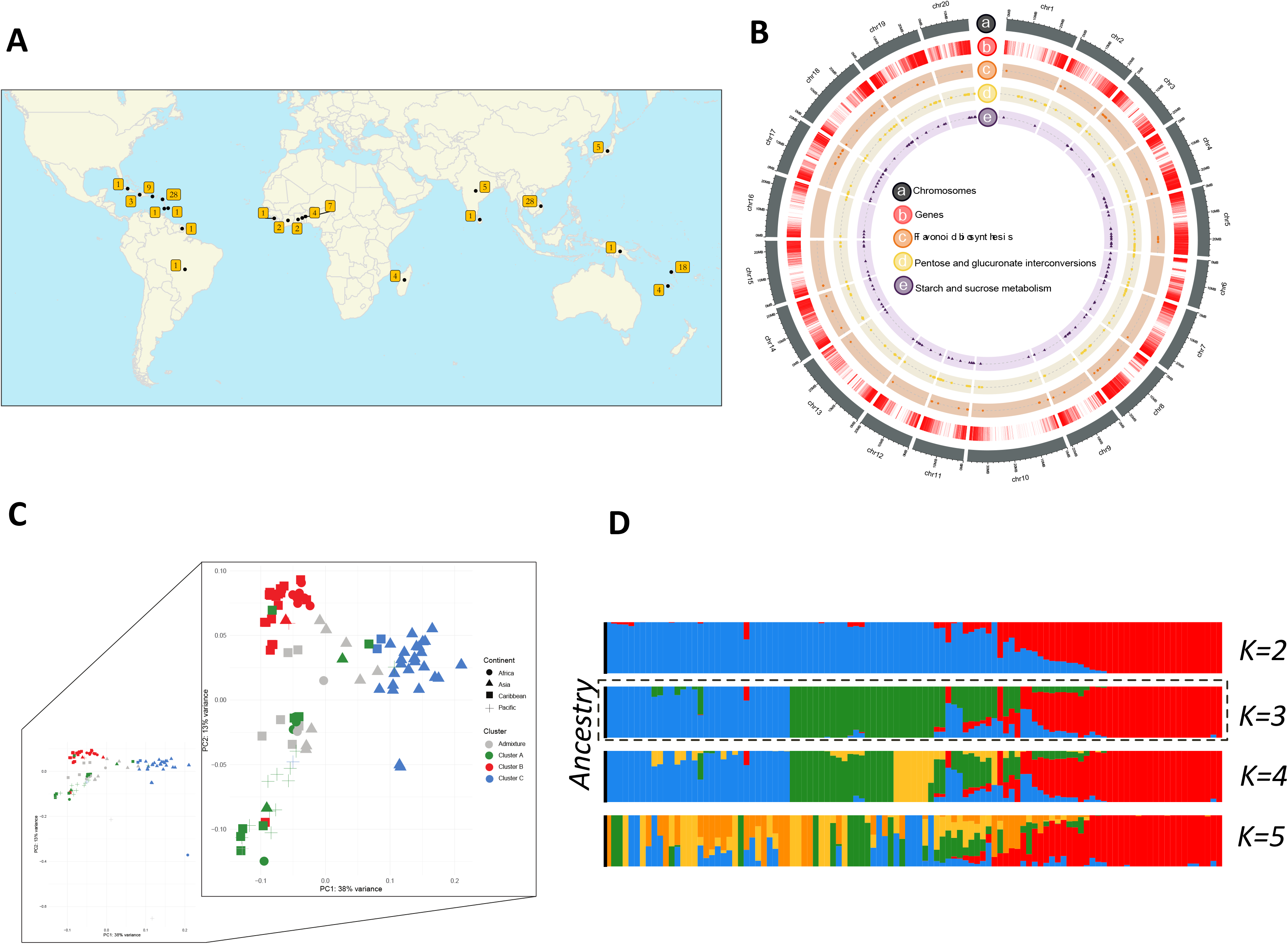
Population structure of *Dioscorea alata*. A: Geographical origin of the 125 *Dioscorea alata* accessions. B: Circos plot of the 20 chromosomes of *D. alata*, the circles represent from outer to the inner circle: chromosomes, gene density, localization of genes from pentose and glucuronate interconversions, genes from starch and sucrose metabolism and genes from flavonoid biosynthesis. C: Principal component analysis depicting the 107 diploid genotypes of *D. alata*. Accessions are coloured according to their assignment to the three genetic clusters after Admixture analysis. The threshold to assign a genotype to a cluster is set at 75 %. D: Admixture barplot showing the distribution of the *K* = 2 to *K =* 4 genetic clusters.

To evaluate the genetic variance among the three clusters, we calculated the fixation index values (*Fst*) for each region in a window of 50kb and a step-window size of 10kb. We found in total 2,141 outlier regions *i.e.,* with significant values of *Fst* (varying from 7.85e^-06^ to 0.64) (Figure 2A, B and C). The comparison of Cluster B *vs* C, showed the highest values of *Fst* and the highest number of outlier regions (881), followed by the comparison of A *vs* C (789) then A *vs* B (471) (Figure 2D). These outlier regions harboured a total of 1,053 unique genes (Figure 2D). We found 11 outlier regions in common among the three genetic clusters, which could be the result of an early purifying selection in *D. alata*. Only four genes were present in these 11 common outlier regions: two closely located genes on chromosome 1 and two closely located genes on chromosome 4. The genes found on chromosome 1 were both annotated as non-specific serine/threonine kinase (EC:2.7.11.1). Similarly, both genes on chromosome 4 are annotated as proteasome endopeptidase complexes (EC:3.4.25.1). Functional analysis of the 1,053 genes based on Gene Ontology (GO) terms showed that various functional pathways were enriched. The most enriched GO terms were tripeptide transmembrane transporter, transmembrane receptor protein binding, stamen filament development, hormone-mediated signalling pathway and lignin biosynthetic process (Figure 2E).

**Figure 2:**
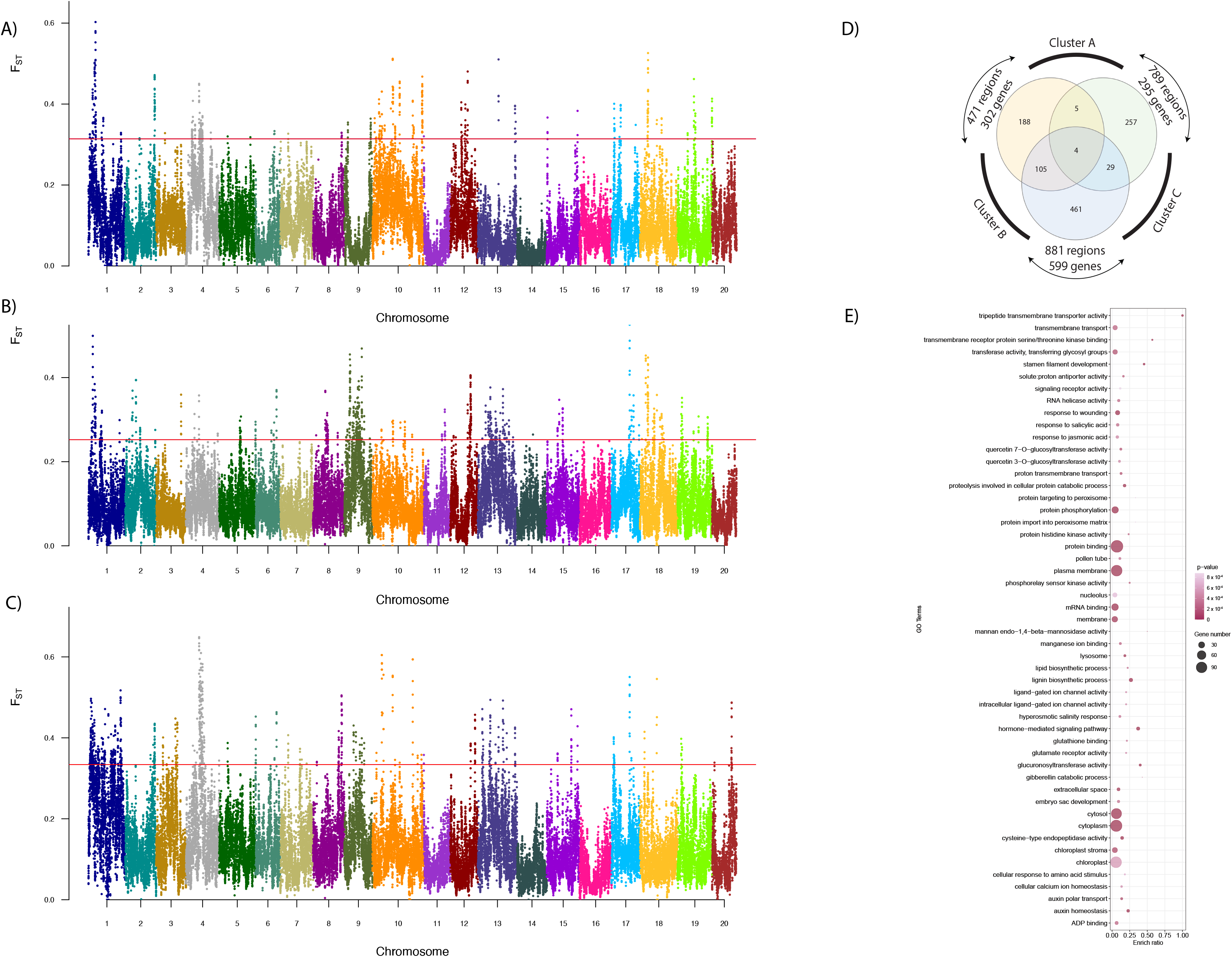
Selection signature found from the comparison of clusters. A: Manhattan plot of Fst analysis of cluster A *versus* B, each colour represents one chromosome. B: *Fst* analysis of clusters A *versus* C. C: *Fst* analysis of clusters B *versus* C. D: Venn diagram of the common genes found among the three clusters, and the number of common outlier regions. E: The gene ontology enrichment of genes found in outlier regions against the whole set of genes.

### Search of genes involved in pathways associated with tuber quality using combined strategies

To better understand the genetic basis of *D. alata* tuber quality, we focused on the genes described to be involved in one of the three pathways targeted (pentose and glucuronate interconversions, starch and sucrose metabolism, and flavonoid biosynthesis). To obtain the most complete set of genes from these pathways, we used two complementary strategies: a keyword search and a comparative genomics approach. For the first strategy, genes annotated on the *D. alata* reference genome (Bredeson et al., 2022) with the enzyme commission numbers (EC number) in one of the three pathways were selected. In total, we could retrieve 322 genes from these pathways encompassing 177 EC numbers (Figure 3A; Supplemental Table 2). The pentose and glucuronate interconversions pathway include most genes (150) and EC numbers (73), followed by starch and sucrose metabolism (136 genes, 76 EC numbers) and flavonoid biosynthesis metabolism (83 genes, 28 EC numbers). To complement the set of annotated genes, a comparative genomics approach was employed using a comparison of 45 diverse plant genomes (Mota et al., 2020) (Supplemental Table 3). Groups of orthologous predicted proteins from different plants (OG), were used to infer the function of proteins that were not annotated by other methods. Since the clustering of proteins in OG does not depend on their previous function annotation but rather on their protein sequence, this method can uncover missing genes from a keyword search. The orthology analysis resulted in a total of 58,916 OG from 1,706,645 proteins (weblink dataverse Cirad). Out of the total proteins analysed, 93.7% could be assigned to an OG. We identified 3,982 OG that are commonly found in all 45 species, despite their significant evolutionary distance. As expected, species that are closely related either phylogenetically or by genus have a greater number of OG in common than distantly ones (Supplemental Figure 4A, B). When we narrowed our comparison to monocotyledons (which comprised 18 species), we found a total of 5,544 OG that are commonly shared among the species. Only three OG were not present in *D. alata* (Figure 3B). The low number of missing groups in *D. alata* species was also observed by Bredeson et al. (2022), when comparing it with species exclusively from the Dioscoreales family.

**Figure 3:**
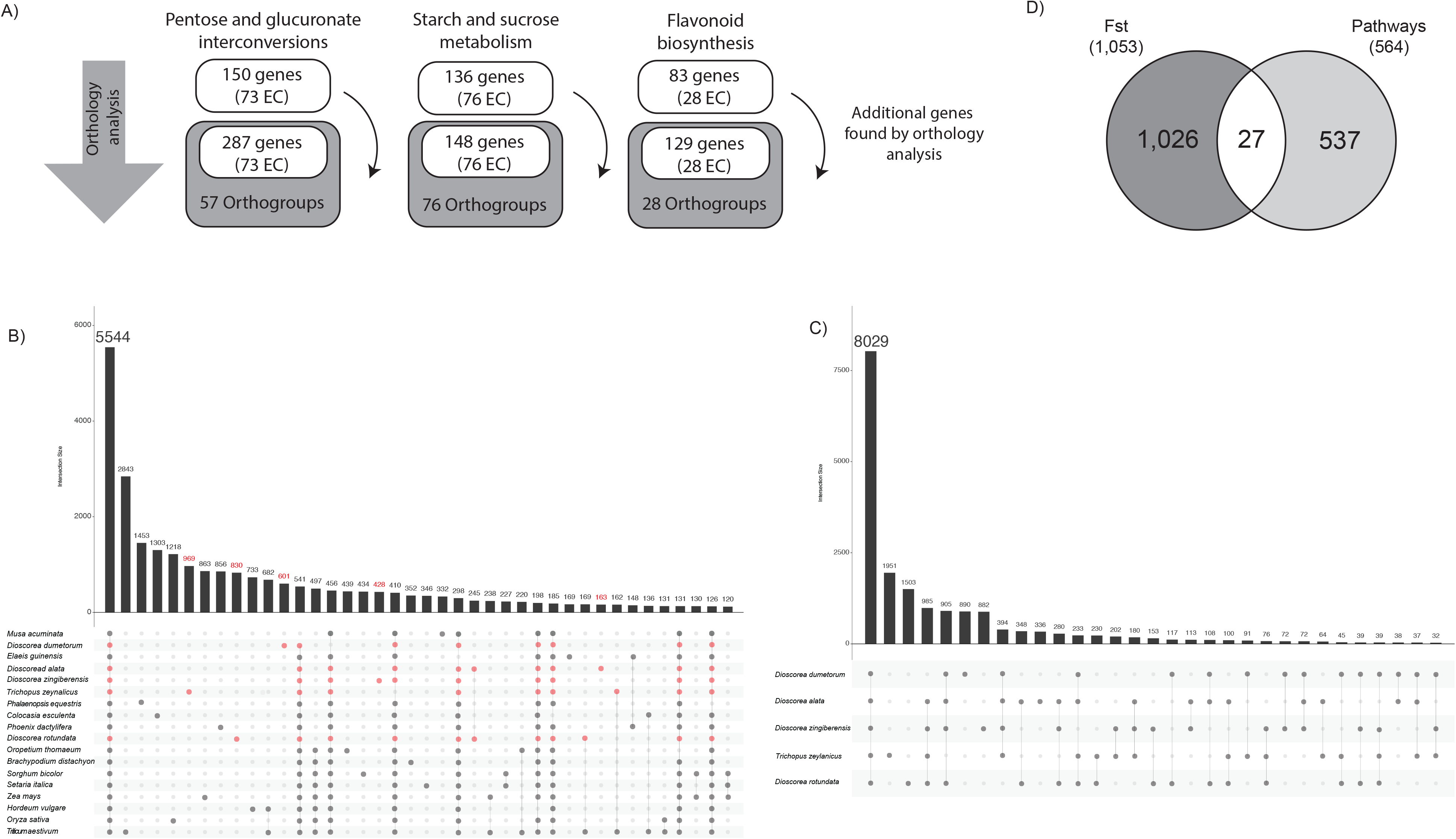
Orthology analysis of 45 plant species. A: Number of genes found in the keyword analysis per pathway and the EC number which they were associated with (light grey). Number of genes found in the orthology analysis, number of EC numbers and number of orthologous groups found (dark grey). B: UpSetR of the comparison of all the monocotyledon plants used in orthology analysis. Each dot represents a connection, and the bar represents the number of orthologous groups found for each association. The red dots show the species from the Dioscoreales family, and the red numbers represent the exclusive associations of Dioscoreales. C: UpSetR of the five Dioscoreales species. D: Intersection between the genes found in the *Fst* analysis and in the pathways.

Among the five Dioscoreales species, we found 8,029 OG in common. *Trichopus zeylanicus* showed the highest number of exclusive OG (1,951) compared to the other *Dioscorea* species (*D. dumetorum*, *D. rotundata*, *D. zingiberensis*, and *D. alata*) and *D. alata* the lowest one (336) (Figure 3C). Additionally, *D. alata* and *D. rotundata* had a high similarity, and likely underwent the same duplication events (Bredeson et al., 2022). We identified 14 OG that comprise 163 exclusive proteins of Dioscoreales (Supplemental Table 4), with most of these proteins (39) belonging to *D. alata*. Based on their EC numbers, most of these OG are hydrolases. They are not exclusive to *Dioscorea* species but their sequence conservation among other species is low, resulting in exclusive OG (*i.e.,* species-specific).

Once we identified all the OG that included representatives from *D. alata*, we narrowed these groups down to the three pathways targeted in this study. For most of the OG found in the three pathways, at least one gene from each of the 45 species was present, or the majority of them (weblink dataverse Cirad; Supplemental Figure 4B). These OG were not exclusive to any particular phylogenetic clade, genus, or species, indicating the importance of these proteins in biological processes for all plant species. Although present in all species, the number of proteins for each OG was uneven, which could be the result of both bias originating from assembly and annotation discrepancies, as well as duplication events that occurred after the differentiation of certain species (in-paralogs) (Pont et al., 2011).

The comparative analysis was not used to increase the number of EC numbers for these pathways but to complete the set of previously annotated genes (Bredeson et al., 2022). We could retrieve new protein coding genes from the comparative genomics analysis which were not found from the keyword search. Thus, we found additional 137 genes for the pentose and glucuronate interconversions pathway, 12 genes for starch and sucrose metabolism pathway and 46 genes for flavonoid biosynthesis metabolism (Figure 3A; Supplemental Figure 5, 6, 7). We were able to associate one orthologous group to each EC number, which contains one or more genes, except for the pentose and glucuronate interconversions pathway, for which the 73 EC numbers were collapsed in only 57 orthogroups. For the other two pathways, each EC number corresponds to an orthologous group. After combining these two approaches, we could retrieve a total of 564 genes from 161 orthogroups for the three pathways, representing 48% more genes than using only one strategy. These genes were distributed along all the chromosomes of *D. alata* with chromosomes 2, 5, 11, and 17 having the highest number of genes and chromosome 16 the lowest (Figure 1B). To determine the polymorphism within the genes found in the three pathways, we selected the open read frame regions since we are only interested in the genic polymorphisms in this study. A total of 406,325 SNPs on 564 candidate genes, from the three pathways was selected. By keeping only SNPs annotated as high impact (*i.e.* non-synonymous, stop codon gain, frame shift) responsible for the most significant differences on the protein sequences, a final number of 4,287 SNPs was used for further analysis, with the pathway of pentose and glucuronate interconversions having the highest number (2,912), followed by starch and sucrose pathway (765) and flavonoid biosynthesis (610).

### Identification of genes under selection in the three targeted pathways

Once we have determined the complete set of *D. alata* genes belonging to the three pathways, we searched for regions under selective pressure, using *Fst* analysis. When comparing the 564 genes from the three pathways to the 1,053 genes from the *Fs*t analysis, we found 27 genes in common including nine genes found exclusively through the comparative genomic approach (Figure 3D, Supplemental table 5). The majority of the 158 SNPs on these 27 candidate genes were on the pentose and glucuronate interconversions pathway (119 polymorphisms in 10 genes) only found between Cluster B (mostly African genotypes) and C (Asian genotypes), followed by starch and sucrose (23 polymorphisms in 7 genes) and flavonoids (16 polymorphisms in 10 genes).

On the pentose and glucuronate interconversions pathway, we found three pectate lyase genes (*Dioal.01G025700.1*, *Dioal.01G025800.1*, *Dioal.01G035100.1*), which together with the polygalactunorase gene (PG) (*Dioal.06G005800.1*) and pectin methylesterase gene (PME) (*Dioal.08G105700.1*) are known as pectin-degrading enzymes involved in the demethylesterification of homogalacturonans (Leng et al., 2017). PME family acts prior to hydrolysis by PG enzyme that degrades the homogalacturonate backbone (polygalacturonate). Pectate lyase gene family is responsible for the cleavage of pectate, which is the product of pectin degradation. From the same pathway, we also found four malectin-like receptor-like kinase genes (MLD) under selection (*Dioal.20G055000.1*, *Dioal.20G055100.1*, *Dioal.20G055200.1*, *Dioal.20G055300.1*). The MLD genes, frequently erroneously described as Receptor-Like Kinases, can bind to pectin, with a preference for more highly methyl esterified pectin, causing a loosening in the cell-wall (Li et al., 2016).

On the starch and sucrose pathway, seven genes were under selection. Among them, two cellulose synthase genes (*Dioal.08G106000* and *Dioal.08G130500*), were under selection between clusters B and C. Additionally, we identified a starch synthase gene (*Dioal.09G032200*) and a nudix-hydrolase (*Dioal.11G000200*), under selection among A and C and among A and B, respectively, a 1,3-beta-glucan synthase (*Dioal.10G014800*), among A and B and a trehalose-6-phosphate phosphatase (*Dioal.04G184200*) under selective pressure among A and C genetic clusters. Those genes are responsible for starch synthesis in plants (Kolbe et al., 2005; Muñoz et al., 2006; Keeling and Myers, 2010). Trehalose phosphate phosphatase is involved in the synthesis of Trehalose consisting of two 1,1-linked glucose molecules (Kolbe et al., 2005). Lastly, we found a beta-amylase gene (*Dioal.04G168200*) under selection. The beta-amylase gene family was previously described to increase the firmness of cooked sweet potatoes when its enzyme activity is low (Banda et al., 2021).

On the flavonoids pathway were found under selection 2-oxoglutarate-dependent dioxygenase (*Dioal.01G016100* and *Dioal.01G016200*) and chalcone isomerase (*Dioal.08G131200.1*). The involvement of these genes was mostly associated with plant-pathogen interaction, and their relationship with quality traits remains to be investigated (Fang et al., 2022).

### Candidate gene association studies for quality traits in *D. alata*

To get a deeper insight in the involvement of the genes annotated from the three metabolic pathways on the tuber quality, we conducted a candidate gene association study. We used phenotypic data collected from two-years field trials on 45 genotypes among the 125 *D. alata* genotypes. They were selected to cover the genetic diversity and to minimize the genetic structure. A set of 10 traits related to starch content, colour indices (Brown Index, the Hue for purple and the Browning over time) and texture were measured. Texture was measured with two different types of probes. A conical probe was used to measure the total area (N sec), which reflects the total force required to penetrate the sample at a constant speed. A flat plate probe was used during a double compression cycle to measure hardness, gumminess, springiness, and cohesiveness. Descriptive statistics of the various tuber quality traits evaluated in this study are shown in Supplemental Table 6. The colour indices BI, HI exhibited wide range of variation (12.57-143 and -11.99-99.92, respectively), showing the presence of varying coloured yam tubers in the association panel. The tuber SC was on average 77.93±2.10 with the minimum (69.46 %) and maximum (81.26 %) observed in CRB417 and Roujol9, respectively. The textural traits also varied extensively within the panel, indicating varying boiled yam qualities of the accessions (Supplemental Table 6). Taking together, our results indicate a wide variation in the tuber quality traits within the 45 genotypes and pinpoint that these traits are largely quantitative in nature.

Based on the general linear model, we obtained significant associations between SNPs and the phenotypic data. We detected a total of 22 quantitative trait nucleotides (QTNs) distributed on 10 chromosomes, for seven out of the ten traits evaluated in this study (Table 1). Hereafter, we present in more details five key genes associated with textural traits, starch content and colour. The expression profiles of the candidate genes were assessed in six phenotypically diverse genotypes using transcriptomic analysis.

Out of the 22 QTNs, 17 were detected for texture traits with fourteen and three within pentose and glucuronate interconversions, and sucrose and starch pathways, respectively (Table 1 and Supplemental Figure 8). Within pentose and glucuronate interconversions pathway, we found pectin esterase (*Dioal.19G187400.1*) and xylose isomerase genes (*Dioal.13G053400*) associated to springiness and gumminess. Leucine-rich repeat (*Dioal.10G024400)*, protein kinases (*Dioal.02G062700*, *Dioal.02G062800*), a malectin-like encoding gene (*Dioal.20G055400*) were associated to springiness and had a stop gain caused by a transition and by a transversion. Two other malectin-like genes found under selection were associated to hardness. Within starch and sucrose pathway we found three candidate genes associated to gumminess, total area and hardness, ɑ-amylase (*Dioal.07G051400.1*), 4-ɑ-glucanotransferase *(Dioal.07G067100)* and beta-glucosidase (*Dioal.13G093500*), respectively.

Pectin methylesterase (*Dioal.04G130600.1*), was associated to gumminess and total area and found under selection. A tuber pectin methyl esterase activity has been identified as a potential factor impacting cooked potato textural properties (Ross et al. 2011). The SNP S4_20715506 (C/T) located in the only exon of *Dioal.04G130600.1* (Figure 4A), results in an amino acid change (H to W), with a potential impact on the enzyme activity. We observed that the C-allele at S4_20715506 decreases gumminess and total area (Figure 4B and C). None of the accessions used for transcriptome analysis had the T-allele, hence we were unable to evaluate the expression pattern of this gene between two haplotypes. Nonetheless, our results suggest that allelic variation in *Dioal.04G130600.1* probably affects the enzyme activity, leading to varying textural properties of boiled yam.

**Figure 4:**
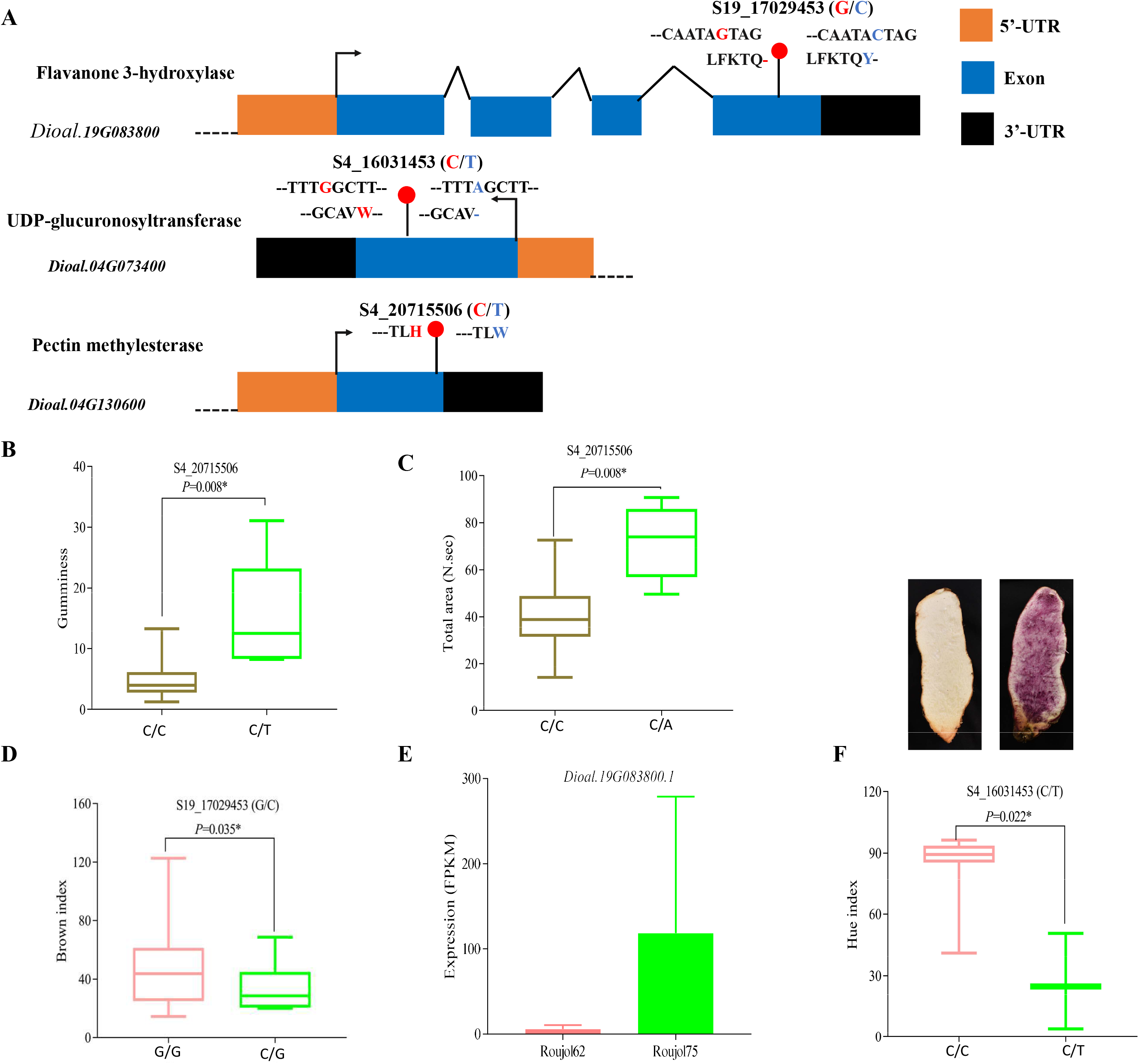
Candidate genes associations and gene expression profile. Selected genes and their expression profile, and allelic effect on colour and texture traits. A: gene structures (*Dioal.19G083800, Dioal.04G073400* and *Dioal.04G130600*) with positions of the significant SNPs, their alleles and resulting amino acid changes; S4_20715506 (C/T) effect on B: gumminess; C: total area; S10_17029453 (G/C) effect on D: brown index (BI); E: Transcriptome based expression of *Dioal.19G083800* between two genotypes with G and C alleles (FPKM= fragments per kilobase of transcript per million fragments mapped); FPKM values was extracted from three biological replicates. S4_16031453 (C/T) effect on F: Hue index leading purple colour change displayed by the two representative tubers. Means were separated by two-tailed *t*-test at 0.05 probability.

Beta-glucosidase, associated to hardness, is a cellulase enzyme playing a part in metabolism of cell wall polysaccharides (Sampedro et al., 2017). The alternative C-allele at the qH13.2 had a strong effect on hardness (62%) by modifying the splice donor on position 837 in the gene *Dioal.13G093500* (Table 1). The genotypes CRB47, Roujol49, and Roujol75 which are homozygous for the G-allele showed an overall higher expression than the genotypes with the C-allele (CRB96, Roujol62, and Roujol9).

The variation of the starch content in the association panel was associated with three QTNs (Table 1). They were annotated as 1,3-beta-glucan-synthase (*Dioal.10G014800*), beta-amylase (*Dioal.15G119500*) and glucose-1-phosphate adenyl transferase (*Dioal.17G112400*). The 1,3-beta-glucan-synthase had the highest effect on the starch content and was also found in an outlier region by our *Fst* analysis (Supplemental Table 5). A recent study demonstrated that manipulation of this gene induces different starch and polysaccharide profiles in barley grain (Garcia-Gimenez et al. 2020). The genotypes harbouring the G-allele had a premature stop codon instead of a glycine. They had 14% less starch content compared to the genotypes with the T-allele. We speculated that the gain of a stop codon results in a non-functional or altered protein which affects the SC.

Lastly, within the flavonoid biosynthesis pathway, we identified two QTNs qHI4.1 and qBI19.1 for Brown index (BI) and HUE index (HI), respectively (Table 1). The gene *Dioal.19G083800.1* encodes for flavanone 3-hydroxylase (F3H). It catalyses the 3-beta-hydroxylation of 2S-flavanones to 2R,3R-dihydroflavonols which are intermediates in the biosynthesis of major pigments (flavonols, anthocyanidins, and proanthocyanidins) (Falcone Ferreyra et al., 2012). *Dioal.19G083800.1* contains SNP S19_17029453 (G/C) in the last exon, with the G-allele leading to a premature stop codon, a truncated protein and likely a non-functional enzyme (Figure 4A). S19_17029453 was significantly associated with BI. The genotypes with the G-allele tend to have brown colour in contrast to the genotypes with the C-allele displaying mostly whitish tuber flesh (Figure 4D). Thus, this mutation may block the formation of flavonoid compounds and redirect the phenylpropanoid pathway to the formation of other brownish compounds like oxidized phenolic acids. Expression pattern of *Dioal.19G083800* between two contrasting accessions with G/C alleles showed that it was expressed nearly 20 times higher in heterozygous (Roujol75; C/G) than in homozygous (Roujol62; G/G) genotypes (Figure 4E). Overall, these results suggest that a functional mutation in *Dioal.19G083800* partly modulates colour formation in yam tuber.

Another key gene detected for colour variation (purple and non-purple yam varieties) was *Dioal.04G073400*, encoding for UDP-glucuronosyltransferase (UGT). Anthocyanins are unstable water-soluble pigments. UGTs are key enzymes that stabilizes anthocyanidin by attaching sugar moieties to the anthocyanin aglycone (Li et al. 2016). In this study, the T-allele at the SNP S4_16031453 leads to a stop codon in the middle of *Dioal.04G073400* (Figure 4A), which will probably result in a non-functional enzyme. Intriguingly, all accessions with the T-allele displayed purple tuber (very low HI score) (Figure 4F). This could be explained by the fact that non-functionality of an enzyme producing uncoloured flavanones could redirect the pathway to anthocyanins biosynthesis resulting in the purple colour of the tubers.

Altogether, the candidate gene association analysis showed that variants in several genes affect the functions and potentially the expression of key structural genes involved in the regulation of yam tuber quality traits. Interestingly, only 38% of the genes found in this analysis were retrieved from the keyword annotation, while the remaining were retrieved by comparative genomics.

## DISCUSSION

In this study, the use of comparative genomics and genome sequence analysis of 127 *Dioscorea alata* accessions enabled the identification of candidate genes in the main metabolic pathways involved in yam tuber quality. The functional annotation of the *D. alata* genome remains suboptimal since there are few genomic data available for this crop (Chaïr et al., 2022). Only recently a complete *D. alata* genome reference was published (Bredeson et al., 2022). Therefore, the comparative genomics combined with keyword search used in this study, allowed the identification of novel genes associated with the three metabolic pathways targeted, pentose and glucuronate interconversions, starch and sucrose metabolism, and flavonoid biosynthesis pathways. We found more paralogs for each metabolic pathway, even if the number of ECs remained the same. We annotated orthologous groups using the conserved domains, which facilitated the correction of incomplete or outdated gene assignments, such as for Malectin-like genes. In addition, based on the orthology analysis we found gene structure errors, such as for OG0082990 and OG0083049 (weblink CIRAD dataverse), which calls for manual curation of important gene families to further improve the functional annotation of *D. alata* genome.

We found that various gene ontology categories were enriched from the genes under selection between the three identified genetic groups. The most enriched pathways under selection were transmembrane transport, protein phosphorylation, response to wounding, hormone signalling, lignin biosynthetic process, known to play important physiological roles in defence response during plant-stress interactions (Zhang and Sun, 2021). This indicates most probably a selection for adaptation to biotic and abiotic conditions. The downsizing to the three pathways revealed under selection cellulose synthase, required for primary and secondary cell wall cellulose synthesis (Richmond and Somerville, 2000), pectate lyase and PG known as pectin degrading enzymes. Additionally, we also identified genes involved in starch synthesis, nudix hydrolase, soluble starch synthase 3 and 1-3 beta-glucan synthase (Miao et al., 2017), confirming the importance of starch content in yam during farmer selection and most probably during domestication (Meyer and Purugganan, 2013). Genes for the pentose and glucuronate interconversion pathway were found only between African and Asian genetic groups. This result deserves further investigation to understand this exclusivity. Five genes from the pectin and starch-sugar pathways were found in common between the *Fst* analysis and candidate gene association study, showing that the quality of yam tubers was also a factor of differentiation and selection during yam evolution.

The candidate gene association analysis enabled the identification of fourteen pectin and three starch candidate genes associated with texture, confirming the involvement of the cell wall and starch content in boiled yam texture (Kouadio et al., 2013). Among the seventeen candidate genes, three have high effect and are involved directly in the cell wall composition. Beta-glucosidase explains more than 60% of the hardness confirming it as a major QTN in boiled yam texture. Its role in cell wall thickness has been demonstrated in different crops such as potato (Ross et al., 2011) or barley in which it participates in endosperm cell wall degradation during germination (Leah et al., 1995). In cotton (*Gossypium hirsutum*), the overexpression of the *GhBG1A* a gene encoding beta-glucosidase repressed fiber length but promoted cellulose biosynthesis resulting in thicker fiber cell wall (Watcharatpong et al., 2020). The four remaining predicted genes with high effect on texture parameters were involved in pectin pathway. Pectin methylesterase, found also under selection, encodes for an enzyme which plays an important role in both pectin remodelling and disassembly and consequently in firming and softening of cell wall (Atmodjo et al., 2013). In our study, it has a negative effect on both total area and gumminess indicating its involvement in pectin degradation during yam boiling, such as in cooked potato (Ross et al., 2011). In cassava, its involvement in the root softening process during cassava retting was demonstrated in addition to pectin/pectate lyase and polygalacturonase genes (Ngolong Ngea et al., 2016). These two last genes were found under selection but not associated with any trait investigated. Finally, Xylose isomerase seems to play an important role in boiled yam texture. It was associated with gumminess and springiness most probably by catalysing the reversible isomerisation of pentoses such as arabinose, one of the components of pectin side chain (Mu et al., 2018).

Such as for cereals and other tuber crops, the total starch content of yams is a primary determinant of tuber quality. Preferred mealy cooking yam cultivars had significantly higher starch content (Kouadio et al., 2013). Among the three genes associated with starch content, the 1,3-beta-glucan synthase had the highest effect. It is often considered to be a cellulase family member, plays an important role in cellulose structure. In this study this gene family was found under selection and predicted to be associated with starch content. The (1,3;1,4)-beta-D-glucans are most abundant in walls of the cereals, specifically in the starchy endosperm of grain, where they can contribute up to 70% by weight of the walls in barley, rye, and oats (Burton and Fincher, 2012). Whether it is related to starch content on yam or other tuberous starchy crops remains unclear.

The colour is a determinant key trait in yam varieties adoption. While in West Africa, white colour is preferred, in the Pacific or Caribbean colourful plates are more appreciated. In our study the flavanone 3-hydroxylase (F3H) was found involved in brown index with a high effect. This gene is involved in the accumulation of catechins in tea plants (*Camelia sinensis* L.) (Singh et al., 2008). In carnation flower (*Dianthus caryophyllus* L.) it has been found involved in colour and fragrance (Zuker et al., 2002). RNAi-Mediated silencing of the F3H confirmed that this gene is one of the key enzymes required for the biosynthesis of flavonoids in strawberry fruit (Jiang et al., 2013). The UDP-glucosyltransferase was found associated with purple colour confirming its role in colour formation in yam tuber. None of these genes were found by Wu et al. (2015) through transcriptome analysis of two contrasting *D. alata* genotypes for tuber purple colour, or by a genome-wide association analysis (Dossa et al., 2023). These results indicate that more genes could be involved in this trait. Moreover, we mainly focused our study on SNPs located in coding regions, and only structural genes involved in three targeted pathways were investigated. This approach therefore excludes the regulatory genes such as transcription factors as well as other candidate genes and genome components that control agronomic traits. Similarly, we did not observe significant gene expression changes between contrasting genotypes for the all identified candidate genes, probably because coding SNPs mainly affect protein functions but at a less extent transcript levels.

In conclusion, quality is one of the main criteria selected during evolution and adaptation, which is reflected in the genes identified in this study. We found twenty-two candidate genes, in pentose and glucuronate, sucrose and starch and flavonoid biosynthesis pathways, associated to the three main attributes of boiled yam: texture, starch content and tuber colour. This is the first work that considers the different cell wall constituents and their effect on texture. In addition, we were able to confirm the expression of five candidate genes in these pathways. Further validation of these results on a larger panel of associations by performing a genome-wide association analysis combined with metabolomic and transcriptomic analyses will confirm the robustness of this approach. Validation of the identified genes will pave the way for favourable allele pyramiding in breeding programmes. Moreover, the use of comparative genomics and the keyword search approach to retrieve genes from the three pathways has proved effective. We have been able to retrieve new protein-coding genes that were not found in the keyword search. This approach, applied to orphan crops for which few genomic resources are available, will improve the annotation of their genomes.

### Material and Methods

#### Plant material

This study used 125 genotypes from *Dioscorea alata* and two outgroup genotypes, belonging to *D. rotundata* and *D. trifida* species, collected in 19 different countries. Genotypes were selected to cover a maximum of worldwide diversity (Figure 1; Supplemental Table 1) based on previous studies (Sharif et al., 2020). These genotypes were used to find genomic polymorphism, compared to the *D. alata* reference genome. From the whole set used in this study, a total of 45 *D. alata* accessions harbouring a high genetic diversity were planted from 2018 to 2020 in May each year and harvested at full maturity in the following year between January and March at three different locations in Guadeloupe (Godet, Roujol and Duclos). The three locations have contrasting pedo-climatic conditions (Cormier et al., 2021). The phenotyping was performed on freshly harvested tubers. Finally, a set of six genotypes was selected from the list, based on their previously described phenotypic traits, to be used for transcriptome sequencing (Supplemental Table 1).

#### Phenotyping for quality traits

Colour was analyzed by computerized image analysis techniques (Detailed method is described in Supplemental Note 1). Images were collected on three tubers per genotype (45) and location (3) during the 2018 cropping season. To test the repeatability and accuracy of the system, a custom colour chart was placed on every picture. Acquired images were analyzed using the Rvision library (Garnier and Muschelli, 2022) with R programming language (R Core Team, 2021). Different colour indexes were tested to characterize purple yam. Finally, the Hue was found to be discriminant within the studied diversity panel. Brown (BI) index was also calculated according to (Buera et al., 1986).

Starch content was predicted using near infrared spectroscopy (NIRS). Two replicates of yam flour samples were scanned with a FOSS-NIRSystems model 6500 scanning monochromator (FOSS-NIRSystems, Silver Spring, MD, USA) equipped with an autocup. The spectroscopic procedures and data recording were conducted with ISIscan (TM) software (FOSS, Hillerød, Denmark). The model was calibrated using 2016 and 2017 data and validated on an external independent 2018 dataset. (Supplemental Note 2).

The texture was measured on steam-cooked yam by penetrometry using the TAX-TPlus texture analyzer (Stable Micro Systems, Ltd., Surrey, UK). Yam tubers harvested at Godet 2019 were used for this analysis. Three tubers per variety were sampled and divided into three equal sections (proximal, distal, central) and prepared as described in Supplemental Note 3. Each part was used to produce three cubes at the central section. Each cube was steam-cooked up to 15 min, followed by a cooling time of 7 min, corresponding to a cube temperature of 45°C. Total area was calculated from the puncture test, while four texture profile analyzer parameters were computed from the force-time curve: hardness (N), cohesiveness, gumminess, and springiness (Supplemental Note 3).

#### Whole-genome sequencing

The genomic DNA from 127 genotypes of *Dioscorea spp*. was extracted from leaves according to a protocol using mixed alkyl trimethylammonium bromide (MATAB) buffer and NucleoMag Plant Kit (Macherey–Nagel, Germany) already described by (Cormier et al., 2019). Sequencing libraries were prepared as described in Dossa et al. (2023). Paired-end high throughput sequencing (2 × 150 bp) was performed on an Illumina NovaSeq 6000 instrument on GeT-PlaGe platform, (Toulouse, France) and Genewiz company (Leipzig, Germany).

The reads quality and mapping and Variant discovery and filtering were performed as described in Dossa et al. in press. For the filtering, the following parameters were used: --max-missing 0.5 --minQ 30 --minDP 10 --maxDP 200. For gene candidate purposes, we selected only the polymorphisms annotated with a high impact on the amino acid sequences.

#### Genetic population analyses

Among all the 127 genotypes of *Dioscorea* spp., 107 diploid genotypes (Supplemental Table 1) were selected for further population genetics analyses. We calculated the population structure using the original filters described above, in addition to: --maf 0.01 --max-alleles 2 --min-alleles 2 --remove-indels --max-missing 0.5 --thin 10 --ld-window 50 --min-r2 0.1. The final VCF file obtained from this analysis was used for population structure analysis using ADMIXTURE version 1.23 (Alexander and Lange, 2011). After testing the replicates of K = 1 to 15, we limited the ancestry threshold to 75%, where genotypes with a value lower than this were considered in the admixture group. After Bayesian clustering analysis, populations were redefined according to the results obtained.

To identify genes under selection, *Fst* analysis was conducted using vcftools with an *Fst*-window-size of 50kb and a *Fst*-window-step of 10kb. A cutoff of the 5% top values and a Weibull distribution analysis on regions with extreme *Fst* between the different genetic clusters identified, were used to select candidates. Subsequently, we searched for the genes on these regions using BEDTools intersect version 2.29 (Quinlan and Hall, 2010).

The gene ontology enrichment analysis of the genes from outlier regions was performed with KOBAS-i webserver (Bu et al., 2021) in order to find out their biological significance.

#### Comparative genomics and orthology analyses

For comparative genomics analyses 45 species from diverse plant lineages were selected (Supplemental Table 2). The species used in this analysis were chosen based on the completeness of their genomes, phylogenetic distribution along the tree of viridiplantae and in the use of plant-models. These plants belong to the monocotyledons and dicotyledons, encompassing 17 different clades. Five genomes belonging to Dioscoreales were added (*Dioscorea alata, D. rotundata, D. dumetorum, D. zingiberensis, Trichopus zeylanicus*). The completeness score of each plant proteome was verified by BUSCO (Simão et al., 2015) using the Viridiplantae dataset of BUSCO. We chose a threshold of 70% of completeness to include public proteomes. For the inference of orthologous groups, OrthoFinder software v2.4.0 (Emms and Kelly, 2015) was used with the default parameters. Gene presence and absence of species in orthogroups was represented by UpSetR (Lex et al., 2014), using a binary matrix as input.

#### Metabolic pathways selection and gene retrieval from D. alata

We downloaded the complete list of enzyme codes (EC) from the pathway of pentose and glucuronate interconversions (map00040), starch and sucrose metabolism (map00500) and flavonoid biosynthesis (map00941) from KEGG database (https://www.genome.jp). These EC numbers were used to retrieve the genes annotated in the GFF file of *D. alata* (https://phytozome-next.jgi.doe.gov/info/Dalata_v2_1). The genes annotated with these EC numbers were further searched in the orthogroups produced by our comparative genomics study, and all the homologous genes from *D. alata* were retrieved. The VCF file was reduced to the middle coding sequences of the annotated genes of interest.

#### Candidate gene association analysis

Three VCF files, one for each target metabolic pathway genes, containing non-synonymous SNPs were extracted from the whole VCF file and further filtered for minor allele frequency ≥ 0.05, missing rate < 20% using TASSEL5.0 (Bradbury et al., 2007). The associations between the polymorphisms from each metabolic pathway candidate genes and the corresponding phenotypic traits were tested using TASSEL5.0 based on the General Linear Model. Only significant variants with P ≤ 0.001 were retained. The association analysis was conducted with phenotypic data from each location independently. The effect of alleles at significant SNPs was assessed by comparing phenotyping data for haplotype groups. A Student’s t-test was used to compare the groups of haplotypes (P<0.05) in the R4.0.23 software with the “ggpubr” packages and “rstatix”

#### Transcriptome analysis of six D. alata genotypes

RNA was extracted from six of the most *D. alata* diverse genotypes, based on phenotypic traits. Three biological replicates for each genotype were used. For each genotype, a 50 mg sample cut from the middle of freshly harvested tuber was ground in liquid nitrogen and total cellular RNA was extracted using a Sigma-Aldrich (St. Louis, MO) Spectrum^™^ Plant Total RNA kit with a DNAse treatment. This was followed by quantification by the Invitrogen (Carlsbad, CA) Quant-iT^™^ RiboGreen® RNA Reagent based on the manufacturer’s protocol and verification of RNA quality by 5200 Fragment Analyzer^™^ System (Agilent, Santa Clara, CA) profile.

Synthesis of cDNA and construction of libraries were done with Illumina, Inc. TruSeq RNA Sample Preparation v2 Kit (San Diego, CA). Fragment size of selected cDNA were between 200pb and 400pb. The 18 libraries were indexed, mixed and sequenced using one lane of an Illumina HiSeq 3000 sequencer with the 2 x 150 cycles, paired-end, indexed protocol (Genewiz facility, Liepzig, Germany).

The quality of the raw reads from the 18 libraries were assessed by FastQC version 0.11.7 (Andrews, 2010). The low-quality sequences and the Illumina adapters were trimmed by Trimmomatic version 0.39 (Bolger et al., 2017). Trimmed reads were quantified using Kallisto version 0.46.1 (Bray et al., 2016) using the reference genome of *D. alata* (Bredeson et al., 2022). To conduct the expression analysis of the genes, we used the log10 of the average value of transcripts per million over the three biological replicates. A functional annotation was performed using the conserved domains of PFAM version 35 and a gene ontology annotation using Pfam2go (http://current.geneontology.org/ontology/external2go/pfam2go).

All the bioinformatics analyses were performed with the support of MESO@LR-Platform at University of Montpellier, CIRAD UMR-AGAP HPC (South Green Platform) and IFB core cluster.

## DATA AVAILABILITY

The Illumina HiSeq 3000 sequencing raw data are available in the NCBI SRA (Sequence Read Archive), and transcriptome data are under the BioProject number: PRJNA918625. The phenotypic datasets are available from the corresponding author upon request.

## ACKNOWLEDGMENTS

We are grateful to Christian Mestres for his valuable advice on yam quality and its attributes. We gratefully acknowledge Nancy Terrier for her critical reading of the manuscript. Our greatest debts go to CRP-RTB, IITA and PRC genebanks and Juliane Kaoh, Mamy Tiana Rajaonah, Senanayake Ravinda Lakshan, and Babil Pachakkil for providing us with plant leaves material. We are also thankful to Elie Nudol, Marie-Claire Gravillon, Christophe Perrot and Erick Maledon for their invaluable help with field work and phenotyping, and Sandrine Causse and Hélène Vignes for laboratory assistance in preparing *D. alata* libraries. Finally, special thanks are due to Sylvain Santoni and Muriel Latreille for their assistance in transcriptome preparation.

---

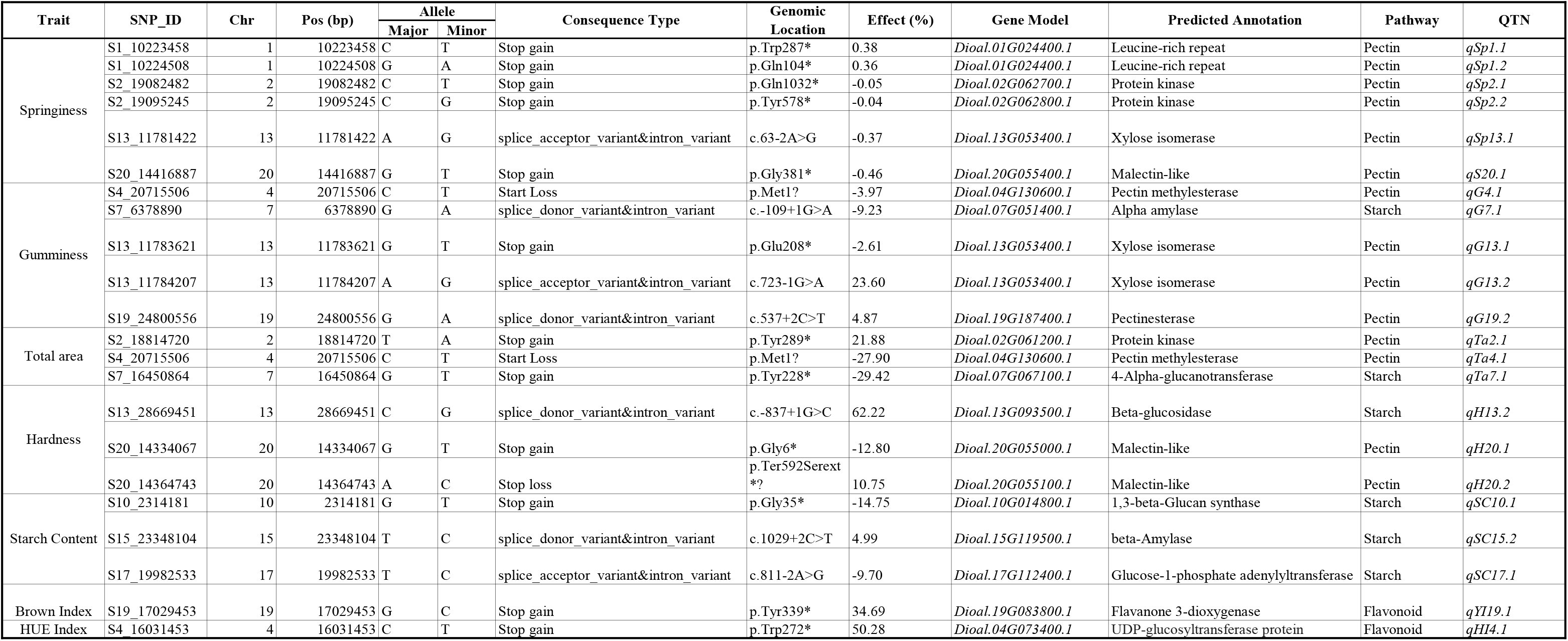

